# A rapid CAT transformation protocol and nuclear transgene expression tools for metabolic engineering in *Cyanidioschyzon merolae* 10D

**DOI:** 10.1101/2024.07.30.605877

**Authors:** Melany Villegas-Valencia, Martha R. Stark, Mark Seger, Gordon B. Wellman, Sebastian Overmans, Peter J. Lammers, Stephen D. Rader, Kyle J. Lauersen

## Abstract

The eukaryotic red alga *Cyanidioschyzon merolae* 10D is an emerging algal host for synthetic biology and metabolic engineering. Its small nuclear genome (16.5 Mb; 4775 genes), low intron content (38), stable transgene expression, and capacity for homologous recombination into its nuclear genome make it ideal for genetic and metabolic engineering endeavors. Here, we present an optimized transformation and selection protocol, which yields single chloramphenicol-resistant transformants in under two weeks. Transformation dynamics and a synthetic modular plasmid toolkit are reported, including several new fluorescent reporters. Techniques for fluorescence reporter imaging and analysis at different scales are presented to facilitate high-throughput screening of *C. merolae* transformants. We use this plasmid toolkit to overexpress the *Ipomoea batatas* isoprene synthase and demonstrate the dynamics of engineered volatile isoprene production during different light regimes using multi-port headspace analysis coupled to parallel photobioreactors. This work seeks to promote *C. merolae* as an algal system for metabolic engineering and future sustainable biotechnological production.

## Introduction

The unicellular red alga *Cyanidioschyzon merolae* 10D was isolated from the volcanic Phlegrean fields near Naples, Italy as a haploid clone without a cell wall (Toda et al., 1995). This alga is considered to represent the simplest free-living eukaryote; composed of a nucleus, a single mitochondrion, one plastid, a simple ER, one Golgi body with two cisternae, few lysosomes, and one peroxisome (Miyagishima & Tanaka, 2021a). *C. merolae* contains a small nuclear genome and had one of the first telomere-telomere complete genome sequences of any model species, with ∼16 Mbp; 4775 protein-coding genes, only 38 introns (Schärfen et al., 2022), and low genetic redundancy (Nozaki et al., 2007). Current genome-wide and multi-omics analysis in *C. merolae* has allowed investigation of the molecular mechanisms related to cell proliferation (Miyagishima et al., 2014; Suzuki et al., 1994), organelle development (Imoto & Yoshida, 2017; Misumi et al., 2005), and various metabolic pathways (Imamura et al., 2010, 2015; Mori et al., 2016; Moriyama, Sakurai, et al., 2014). Genetic manipulation of *C. merolae* 10D has been reported, including precise integration of transgenes in both nuclear (Fujiwara et al., 2019; Minoda et al., 2004; Sumiya et al., 2015) and chloroplast (Zienkiewicz et al., 2017) genomes via homologous recombination (HR) using 200–500 bp targeting sequences (Fujiwara et al., 2017; Takemura, Imamura, et al., 2019). In addition, transgenes are stably expressed, and there appears to be no endogenous silencing of transgenes (Watanabe et al., 2014; Zienkiewicz et al., 2019).

The biomass of *C. merolae* 10D is protein-rich (Villegas-Valencia et al., 2023) and contains valuable bioproducts like heat-stable phycocyanin, carotenoids, and β-glucan (Lang et al., 2022; Rahman et al., 2017; Villegas-Valencia et al., 2023). Because the cells lack a rigid cell wall and are easily disrupted, the introduction of DNA and extraction of intracellular contents is straightforward (Miyagishima & Tanaka, 2021b). The *C. merolae* 10D strain can be cultivated in acidified medium (pH 1–3) and elevated temperatures (42-46 °C) in freshwater and seawater to high cell densities in both lab-scale and outdoor conditions with minimal contamination (Hirooka et al., 2020; Villegas-Valencia et al., 2023). These features suggest that *C. merolae* could be a valuable and scalable platform for metabolic engineering or other recombinant bioproduct accumulation.

Further advances in metabolically engineering this alga are of interest to add value to the algal biomass. There is a growing number of engineered traits in *C. merolae*, including the enhanced generation of triacylglycerol (TAG) without growth inhibition (Sumiya et al., 2015), the incorporation of a plasma membrane sugar transporter from its relative *Galdieria sulphuraria* to enable heterotrophic growth in the presence of exogenous glucose in the dark (Fujiwara et al., 2019), and recently, our demonstration of the production of the non-native ketocarotenoids canthaxanthin and astaxanthin (Seger et al., 2023). Although various genetic tools have been developed in *C. merolae* 10D, current methods for its transformation and standard screening of transformants can take several weeks (Fujiwara et al., 2017; Fujiwara & Ohnuma, 2017; Zienkiewicz et al., 2019). Metabolic engineering attempts in this host are still in their infancy and the development of techniques to achieve mature genetic engineering concepts are still needed. This work seeks to create a user-friendly and standardized transformation approach and we present new modular genetic tools for *C. merolae* nuclear genome transgene expression for metabolic engineering and recombinant product accumulation.

Here, we report an optimized transformation and selection protocol for *C. merolae* 10D, which yields single chloramphenicol-resistant colonies in under two weeks and enables phenotypic screening within three. We present an *in silico* designed and synthetically constructed modular plasmid toolkit which includes several fluorescent reporters. We show its transformation, the metrics of successful homologous recombination, and techniques for fluorescence imaging for the high-throughput screening of transformants. The transgene expression dynamics from the nuclear genome including total soluble recombinant protein accumulation and a first demonstration of metabolic engineering volatile isoprene production from the alga are also presented. This work will serve as a foundation of open-source molecular tools for the *C. merolae* research community and indicates it could be a reliable chassis for further engineering concepts.

## 2. Materials and Methods

### 2.1. Algae culture

*C. merolae* 10D strain (wild-type; NIES-3377) was obtained from the National Institute of Environmental Studies’ microbial culture collection in Japan. *C. merolae* culture and its transgenic lines were routinely maintained in MA2G medium, a modified version of MA2 (Minoda et al., 2004). This medium is a version of Modified Allen’s medium (MA, (Minoda et al., 2004)) wherein several components have doubled concentrations. MA2G consists of 40 mM (NH_4_)_2_SO_4_, 8.0 mM KH_2_PO_4_, 4.0 mM MgSO_4,_ 1.0 mM CaCl_2_.2H_2_O, 100 μM FeCl_3_.6H_2_O, 80 μM Na_2_EDTA.2H_2_O, 92.2 μM H_3_BO_3,_ 18.2 μM MnCl_2_-4H_2_O, 1.54 μM ZnCl_2_, 3.2 μM Na_2_MoO_4_.2H_2_O, 0.34 μM CoCl_2_.6H_2_O, 0.64 μM CuCl_2_-2H_2_O, and 50 mM glycerol. H_2_SO_4_ was used to adjust the pH to 2.5. Glycerol was added as it encourages slightly faster growth of the Cyanidiophyceae and has been shown to encourage respiration (Moriyama et al., 2015). Working stocks of cultures were maintained on corn starch beds on MA2G Gellan gum plates, and 50 mL liquid cultures were shaken at 100 rpm in 250 mL Erlenmeyer flasks with 0.22 μm filter vented caps to allow gas exchange. Cultures were maintained under continuous illumination (90–130 μE) at 42 °C in a Percival incubator (AL-30, Percival Scientific) supplemented with 2– 4% CO_2_. Wild-type and transgenic cultures were cryopreserved in 8% DMSO and stored at −80 °C for long-term preservation.

### 2.2. Plasmid construction

Plasmids were designed to integrate the selectable marker chloramphenicol acetyltransferase (CAT), a fluorophore (one of mTagBFP2 (Subach et al., 2011), mTFP1 (Ai et al., 2006), Clover (Lam et al., 2012), mVenus (Kremers et al., 2006), LSSmOrange (Tsutsui et al., 2008), mKOk (Shcherbakova et al., 2012), mScarlet (Bindels et al., 2017)), and isoprene synthase (IspS) into the intergenic region between the nuclear glycogen phosphorylase (CMD184C) and TATA-box binding protein-associated factor 13 (CMD185C) genes on *C. merolae* chromosome 4 (“HR-L” and “HR-R” arms in Figures 1a, 2b, 3a and 4a) via homologous recombination (HR (Fujiwara et al., 2013)). Gene elements used in our *in silico* genetic designs are listed in Supplemental File S1 and all plasmids can be obtained from the authors or with permission from Genscript.

**Figure 1.**
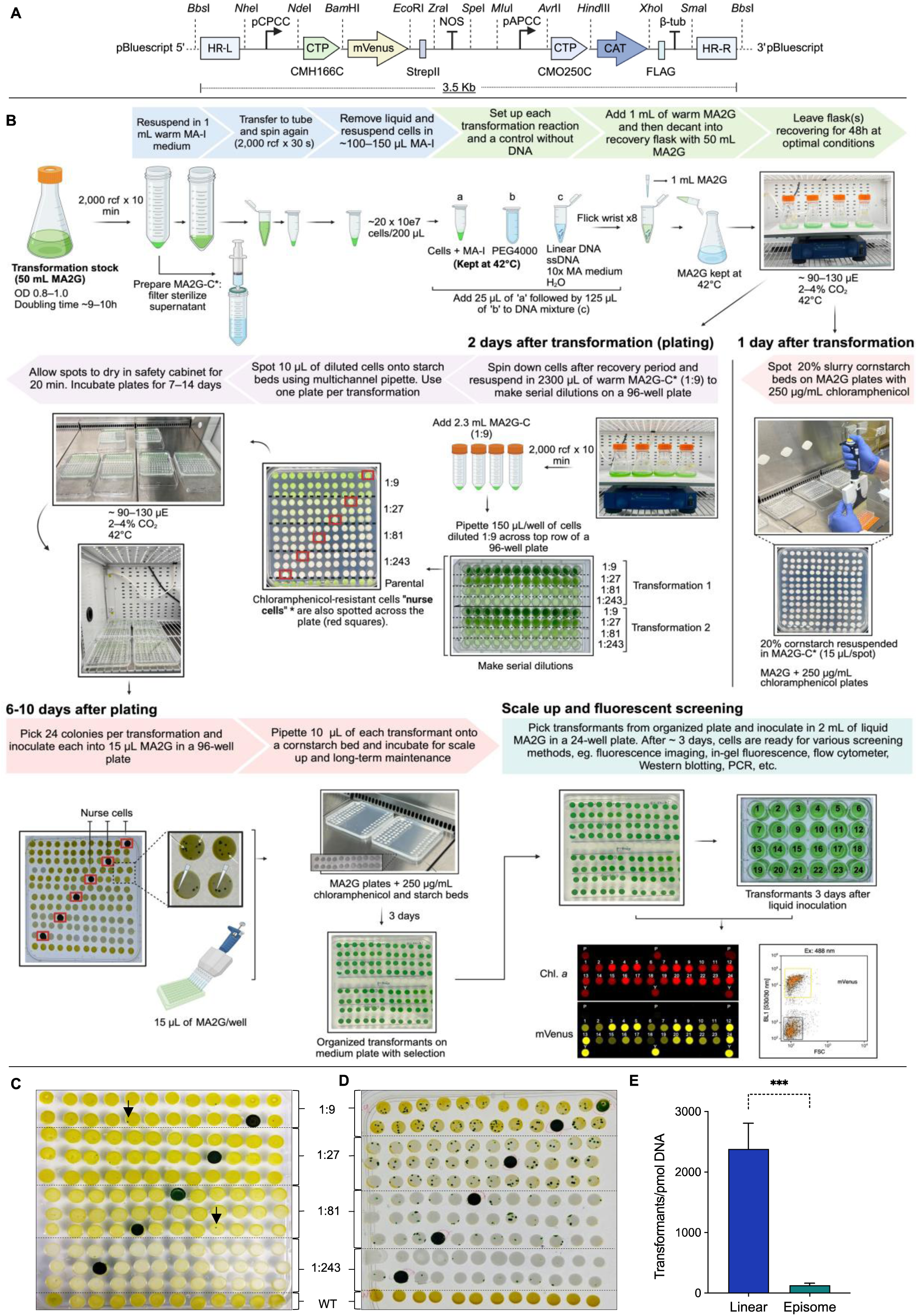
Plasmid design, PEG4000-mediated *C. merolae* 10D transformation and transformation efficiencies. **A –** Synthetic plasmids were designed *in silico* and constructed *de novo* for integration into the 184C-185C neutral site on *C. merolae* chromosome 4. The expression cassette was designed to be modular, with a separate gene of interest and a selectable marker. Each element was separated by unique restriction endonuclease sites as illustrated. pCPCC – phycocyanin-associated rod linker protein promoter, CTP CMH166C – DNA Gyrase B chloroplast targeting peptide, mVenus – yellow fluorescent protein reporter, StrepII – C-terminal peptide tag with stop codon, NOS – nopaline synthase terminator, pAPCC – allophycocyanin-associated rod linker protein promoter, CTP CMO250C – allophycocyanin-associated rod linker protein chloroplast targeting peptide, CAT – chloramphenicol acetyltransferase, FLAG – peptide tag with stop codon, β-tub – *C. merolae* β-tubulin terminator CMN263C. **B** – Workflow of the optimized transformation protocol in *C. merolae* 10D to obtain positive transformants ∼1.5 weeks after selective agent plating. **C+D** – Circular vs. linear DNA transformation. Representative transformation plates ∼ 10 days after inoculating transformants in cornstarch beds. A dilution series of cells transformed with circular (C) and linearized (D) DNA were spotted along with untransformed wild-type cells (WT; last row) and nurse cells (see methods). **E** – Transformation efficiencies compared between linear to circular DNA. *** indicates results of Student’s *t-*test were significantly different (*p* < 0.0001) when comparing the mean number of transformants between linear and episomal transformation.

Endogenous elements such as promoters, terminators, and homology arms were taken from the reference genome of *C. merolae* 10D (Fujiwara et al., 2013, 2017, 2019; Moriyama, Tajima, et al., 2014). The *Staphylococcus aureus* chloramphenicol acetyltransferase (CAT) (NCBI: M58516.1 (Schwarz & Cardoso, 1991)) was used as a selection marker and *Ipomoea batatas* isoprene synthase (*Ib*IspS, NCBI: AZW07551.1 (Ilmén et al., 2015)) for isoprene production. Fluorophore sequences were taken from the sources indicated in Figure 3b. Codon optimization after back translation of amino acid sequences was conducted for the most frequent codon usage for the algal nuclear genome using *C. merolae’s* codon usage table found in the Kazusa database (https://www.kazusa.or.jp/codon/cgi-bin/showcodon.cgi?species=280699). For all genes, native targeting peptides were removed from the amino acid sequence before optimization. Gene synthesis and subcloning were carried out by Genscript (Piscataway, NJ, USA), and plasmids were delivered as lyophilized DNA, transformed, and preserved in *Escherichia coli* DH5α with ampicillin as the selection agent in lysogeny broth (LB). Full annotated sequences of all plasmid elements can be found in Supplemental File S2.

### 2.3. *C. merolae* 10D transformation

The transformation procedure was similar to the ones reported before (Fujiwara et al., 2017; Ohnuma et al., 2008, 2014) with slight modifications, as follows: 1 pmol of linear DNA was used in transformations, which was prepared by PCR amplification using primers targeting the homologous arms with a high-fidelity polymerase (Figure 2A, primer set 1). PCR products (50 μL reactions) were purified using a PCR clean-up kit (ZR-96 DNA Clean & Concentrator) and re-suspended in DNase and RNase-free water. DNA concentration and purity were measured on a NanoDrop One spectrophotometer (Thermo Fisher Scientific, UK).

**Figure 2.**
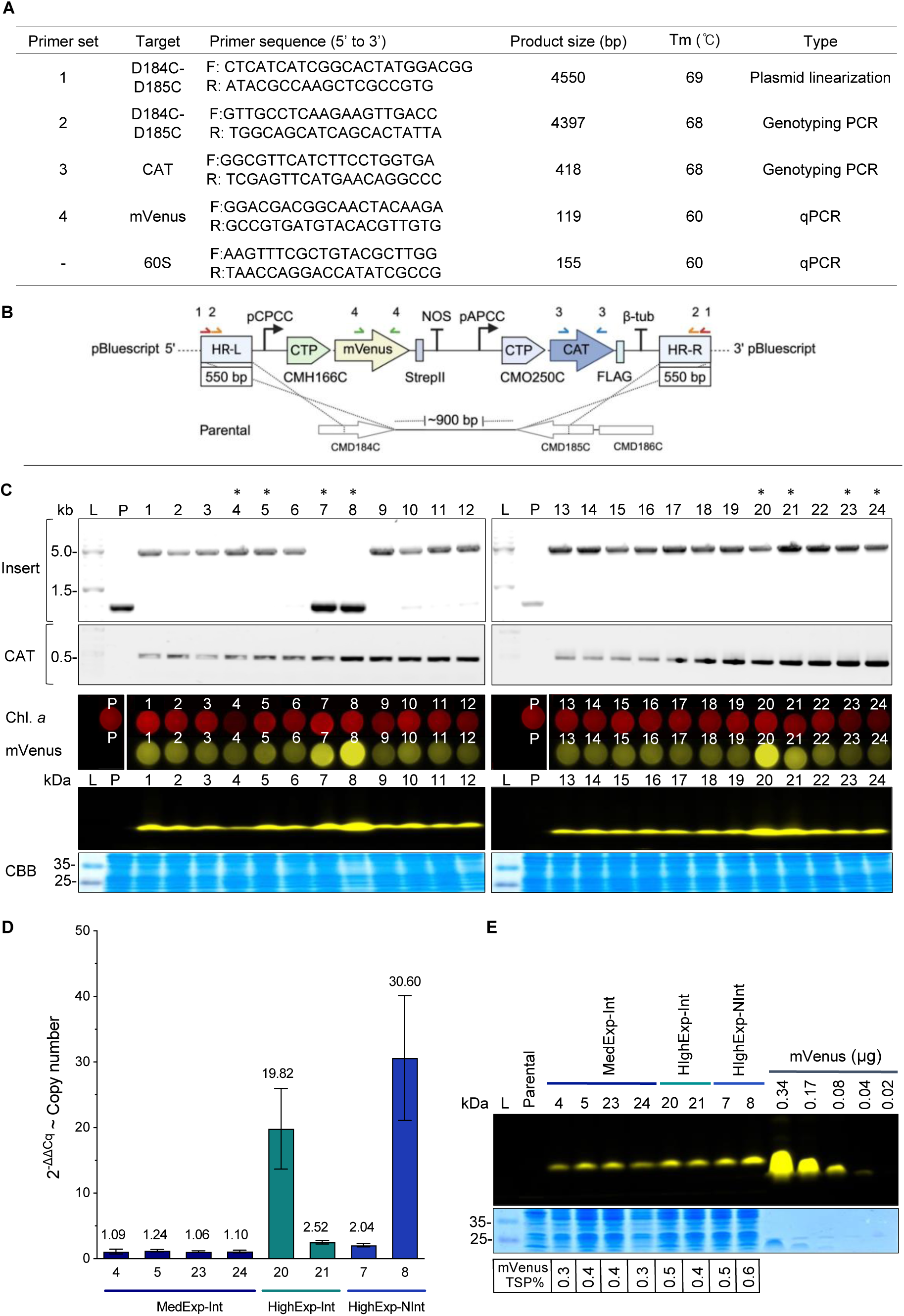
Molecular screening of mVenus-expressing transformants. **A** – Primer sets used in this study to linearize all plasmids before transformation (1), screen for integration at the desired locus and presence of selectable marker (2 and 3, respectively), and estimate the copy number of the gene of interest (4) by quantitative PCR (qPCR). **B** – Schematic diagram of mVenus and CAT insertion into the neutral site between CMD184C and CMD185C by homologous recombination. Introduced linear DNA vector (top) and the genomic structure of the parental (P) wild-type (WT) strain (bottom). Arrowheads indicate the position of the primers used to screen for total insert (∼ 5.0 kb; orange) and the insertion of the CAT gene (∼ 0.5 kb; blue), as shown in C. **C** – Polymerase chain reaction (PCR) confirmation of linearized plasmid integration at the D184-185 neutral locus (insert) and presence of transgene (CAT) in 24 transformants. The predicted size of the parental strain (or off-target insertion) and correctly integrated PCR products is 0.9 kb and 4.4 kb, respectively. The asterisks (*) above the lane numbers show the selected transformants used in further qPCR and TSP analysis. PCR was followed by plate-level fluorescence analysis of the mVenus transformants on Gellan gum plates stained with amido black. Chlorophyll *a* emission (red) and mVenus fluorescent signal (yellow), as well as in-gel fluorescence, were measured to assess the expression level of mVenus in 24 transformants. The expected molecular weight of mVenus is ∼26 kDa. CBB: Coomassie brilliant blue stain is included as a loading control. L: DNA / protein ladder, P: parental strain. Full-length images can be found in Supplemental Figure 1. **D** – Quantitative-PCR analysis of selected transformants to estimate the copy number of the mVenus gene. Transformants 4, 5, 23, and 24 exhibit medium-mVenus-expression with correct integration at locus (MedExp-Int); 20 and 21: high-mVenus-expression, correct integration at locus (HighExp-Int); 7 and 8: high-mVenus-expression and integration outside of target locus (HighExp-NInt). The value of each transformant was normalized against the value from the 60S rDNA gene reference gene to estimate their copy number (n=3). **E** – Percent of total soluble protein (%TSP) analysis of recombinant mVenus expression in transformants measured by in-gel fluorescence. The cell number of the transformants was normalized before extraction and gel loading. A dilution series of purified mVenus from *E. coli* was included for semiquantitative assessment. CBB is shown as a loading control.

To prepare for transformation, cell cultures were maintained in MA2G (MA2 + 50 mM glycerol) medium under continuous light at 90–130 μE, 42 °C and constantly supplemented with 2–4% CO_2_. The doubling time under these conditions is ∼9–10 h for wild-type cells. Wild-type cell cultures were diluted 2–3 days before transformation so that a culture of actively dividing cells with an OD_740nm_ lower or equal to 3.0 was ready prior to transformation (“transformation stock”). One day before transformation, the stock culture was diluted to an OD_740nm_ of ∼0.2 in 50 mL MA2G and grown under the same conditions as above until the OD_740nm_ of the culture reached ∼0.8–1.0. On the day of transformation, polyethylene glycol (PEG)-4000 was prepared (60% w/v) by combining 0.9 g PEG4000 (A16151.30, ThermoFisher Scientific) and 750 μL MA-I (note: same concentrations of MA2 except 20 mM (NH_4_)_2_SO_4_, 2 mM MgSO_4_ + metals, pH adjusted to 2.5) in a 2 mL Eppendorf tube and dissolved at 42 °C with occasional inversion. Next, 50 mL of transformation culture was centrifuged (10 min at 2000 x g), the supernatant filter sterilized to make conditioned media (‘MA2G-C’), and cells washed with 1 mL of MA-I medium kept at 42 °C, and then transferred to a 1.5 mL tube, centrifuged again (1 min at 2000 x *g*) and resuspended in ∼100-150 μL warm MA-I to a total final volume of 200 μL (250x concentrated). In a 1.5 mL tube, ∼1 pmol of linear-(PCR product) or circular-DNA was diluted in water to a total volume of 84 μL and combined with 10 μL 10XMA medium (to bring the transformation mixture to 1XMA-I concentration), and 6 μL 10 mg/mL UltraPure Salmon Sperm DNA Solution (Invitrogen, USA). Salmon sperm DNA was denatured by heating to 98 °C for 5 min and then rapidly cooled on ice before adding. Next, 25 μL of concentrated cells were added to the DNA mixture. A no-DNA control reaction was also prepared as a negative control. Then, one sample at a time, 125 μL of PEG4000 solution was added to the reaction and mixed quickly by flicking the wrist 8-10 times so that the PEG and the cells/DNA were completely mixed. 1 mL of warm MA2G was immediately added to the tube, which was then poured into 50 mL of warm MA2G in a 250 mL vented flask and allowed to recover while shaking at 100 rpm for 48 h under the same cultivation conditions as described above.

After recovery, transformed cells were collected by centrifugation and resuspended in 2.3 mL MA2G-C, a medium in which cells have previously been grown to an OD_740nm_ 0.8-1.0, and has been filter sterilized. MA2G-C was used to dilute cornstarch and cells for plating after transformations. This medium was prepared by growing wild-type *C. merolae* cells in MA2G to a culture OD_740nm_ ∼1.0, centrifuging the cultures, and using the 0.2 μm filtered supernatant. We have observed this medium from *C. merolae* cells in the exponential growth phase accelerates the formation of single colonies on starch beds. Cell suspensions were then serially diluted (1:9, 1:27, 1:81, and 1:243) in a 96-well plate and 10 µL pipetted onto freshly prepared starch beds on 0.77X MA2G Gellan gum plates (0.46% Gellan gum; 120mm x 120mm x 17 mm square petri dishes) containing 250 μg/mL chloramphenicol. Based on our observations, colonies tend to come up faster on 0.77X MA2G Gellan gum plates than on 1X. This dilution series should result in at least one series of starch beds that contains uncrowded colonies appropriate for isolation (∼1-10/spot) across a wide range of transformation efficiencies. The plates are prepared the day before use with 20% (v/v) cornstarch slurry beds (Kobayashi et al., 2010), as shown in Figure 1B. Before spotting the starch, MA2G Gellan gum plates were left open in the incubator for 20 min so that spots that do not run into each other are formed. Briefly, 20% cornstarch slurry was prepared from cornstarch that was washed 3X with sterile H_2_O, resuspended to 50% (v/v) in 75% EtOH, and kept at 4 °C until further processing. For plating, 20% cornstarch was prepared by taking 10 mL from the 50% cornstarch/EtOH stock, centrifuging briefly, washing the pellet 3X with MA2G-C medium, then resuspending to a final volume of 25 mL in MA2G-C. This was poured into a Universal Reagent Reservoir and manually agitated to avoid settling of the starch while pipetting 15 μL spots onto the plates using a multichannel pipette. Chloramphenicol was only added to the Gellan gum + MA2G solid medium, and the cornstarch was antibiotic-free.

Plates with approximately 144 cornstarch spots were inoculated with serially diluted transformants (10 μL per spot), along with 6 spots of nurse cells (Kobayashi et al., 2010) throughout the plate (chloramphenicol resistant and actively dividing cells that may encourage neighboring colony growth; Figure 1B). Nurse cells are spotted on top of a few transformants for ease of plating with the multichannel pipette. Spots were allowed to dry and plates were incubated in CO_2_-supplied Percival incubators under previously described conditions until colony formation appeared ∼6–10 d after plating. Colonies were then picked and resuspended in 15 μL MA2G, and this liquid was used to re-inoculate cornstarch beds for long-term storage and maintenance. Picked liquid samples can also be grown for ∼3–4 d (Figure 1b) and used in further analysis. Isolates were then scaled up in 1 mL of liquid MA2G medium in 24-well plates until dense (OD_740nm_ ∼1.0; after 3 d), and used in plate-level fluorescence, in-gel fluorescence, and flow cytometry assays to confirm the expression of fluorescent reporters.

### 2.4. Transformation efficiencies

Constructs containing mVenus and CAT were transformed into *C. merolae* to determine transformation efficiencies of integrated linear DNA and circular plasmids. The DNA concentration of each was normalized to 1 pmol. Transformations were carried out in triplicate as previously described. 10 days after plating, colony-forming units (cfu) were counted using images of the plates and a software for biological-image analysis (Fiji; Schindelin et al., 2012). Transformation efficiencies were calculated based on cfu from the dilution series on each plate (Supplemental File S3), using the following formula:

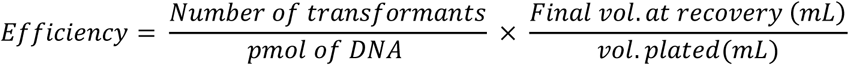

### 2.5. DNA extraction and molecular screening

*C. merolae* strains were harvested at mid-log phase by centrifugation (10 min at 2000 x *g*) and total genomic DNA was extracted from algal pellets using a Quick-DNA Fungal/Bacterial Miniprep Kit (Zymo Research, USA) and a FastPrep-24 5 g bead beating grinder and lysis system (MP Biomedicals, USA) according to the manufacturer’s protocol. DNA extracts were quantified using a NanoDrop One spectrophotometer (Thermo Fisher Scientific, UK).

Q5 High-Fidelity DNA Polymerase (New England BioLabs, UK) and Phusion Green Hot Start II High-Fidelity PCR Master Mix (Thermo Fisher Scientific, Lithuania) were used for PCR according to the manufacturer’s protocols (Table 1). All primers used in this study were synthesized by IDT (Integrated DNA Technologies Inc., Belgium). Primer set 1 was used to linearize target constructs from plasmids before transformation, whereas primer sets 2 and 3 were used to screen transformants for the presence of the insert at the target neutral site and to verify the positive integration of the selectable marker (CAT), respectively.

Quantitative PCR (qPCR) was carried out using a CFX96 Real-Time PCR Detection System to determine the copy number of the integrated mVenus. The copy number of heterologous mVenus in each strain was normalized with that of native 60S rDNA (NCBI: 16997147) as the reference gene. To determine the optimal annealing temperature of the primers, a thermal gradient across 7 different temperatures ranging from 59 °C to 69 °C was used for each primer set in a T100 Thermal Cycler (Bio-Rad). 5 ng of DNA was added to each well in 20 μL reactions and a no template control (NTC) consisting of H_2_O was also included. SsoAdvanced Universal SYBR Green Supermix (Bio-Rad) was used for qPCR according to the manufacturer’s protocol. Standard curves were constructed using serially diluted DNA (1/5, 1/25, 1/125, 1/625, and 1/3125; Supplemental Figure 3) isolated from a mVenus-expressing *C. merolae* strain and the relevant set of primers. For each primer pair, technical triplicates were done for each dilution factor. An NTC was also included for each primer set. PCR and qPCR conditions are shown in Supplemental File S4.

In addition, a semi-quantitative assessment of mVenus accumulation was carried out in selected transformants. The total soluble protein (TSP) fraction of the algal cells was quantified by the bicinchoninic acid (BCA) assay using bovine serum albumin (BSA) as a standard of known concentrations. Samples of TSP extracted from mVenus-expressing transformants, along with a dilution series of *E. coli* produced and StrepTrap (Cytiva StrepTrap XT, Sigma-Aldrich, Germany) column chromatography purified mVenus were analyzed under non-denaturing sodium dodecyl sulfate-polyacrylamide gel electrophoresis (SDS-PAGE) conditions (5% stacking, 15% resolving), followed by fluorescence imaging using excitation wavelengths for mVenus (Figure 2E).

### 2.6. Fluorescence imaging

*C. merolae* transformants cultivated in 24-well plates in 1 mL MA2G were prepared for screening analysis during the mid-log phase. The presence of fluorescent proteins in selected transformants was observed using a ChemStudio PLUS (Analytik Jena, USA) gel documentation system with an eLite xenon lamp and filter wheel extension as previously described (Gutiérrez et al., 2022). Different filters with specific bandpass ranges that allowed the selective excitation and emission of different fluorophores were employed. Plate-level fluorescence was carried out by spotting 10 μL of selected transformants onto a Gellan gum plate with amido black (150 mg/L). The latter was employed to reduce background fluorescence (Wichmann et al., 2018) and it was added to the Gellan gum-MA2 mix by dissolving into the medium before autoclaving. The excitation/emission filters used for each fluorophore are stated in Figure 3D.

**Figure 3.**
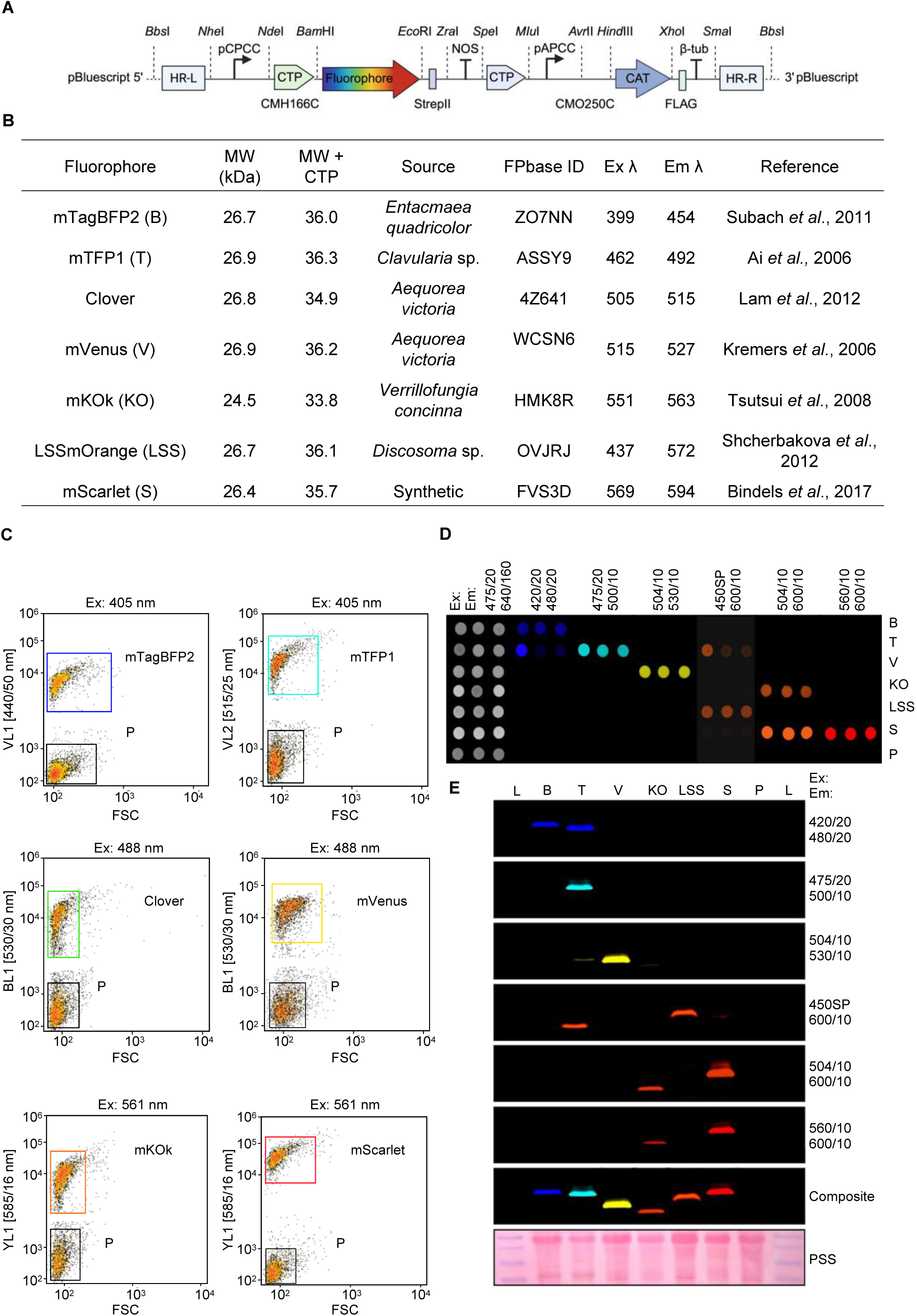
Fluorescent screening of *C. merolae* strains expressing various fluorescent reporters. **A** – Transgenes were codon optimized for *C. merolae* nuclear genome expression and subcloned into a two-cassette system as illustrated. **B** – Fluorescent proteins highlighted in this work, molecular weight of the protein alone or fused to a chloroplast targeting peptide, biological source or synthetic status, Fluorescent Protein Database ID (https://www.fpbase.org), optical properties and sequence source. **C** – Flow cytometry of cells expressing fluorescent reporters that can be separated from the parental strain. mTagBFP2 and mTFP1 signals compared to wild-type (WT) with violet laser (405 nm) excitation and emission from the VL1 and VL2 filter plotted against forward scatter (FSC). Clover and mVenus signals compared to WT were obtained using blue laser excitation at 488 nm and measuring the emission through the BL1 filter, with the populations plotted against FSC. In addition, mKOk and mScarlet signals compared to WT were acquired by using a 561 nm laser for excitation and then visualized with the YL1 emission filter plotted against FSC. **D** – Transformant screening by plate-level fluorescence analysis on Gellan gum MA2 plates stained with amido-black. 10 μL of three representative transformants from each fluorescent strain were spotted horizontally. Signals from each row after emission imaging are shown for each strain along with their respective excitation (Ex) and emission (Em) filter combination sets. Chlorophyll *a* emission (left side). A composite of all images is shown for comparison. **E** – In-gel fluorescence screening of reporter protein signals. One representative transformant from each strain was subjected to SDS-PAGE and imaged using the ChemStudio Plus with the same filter sets as above. The signal from each reporter appears as a single band, which were then combined to create a composite image. Colors in D and E have been incorporated digitally using the VisionWorks ver. 9.0 software. Full-length images can be found in Supplemental Figure 2. MW – molecular weight, MW + CTP – total molecular weight with chloroplast targeting peptide, L – ladder, B – mTagBFP2, T – mTFP1, V – mVenus, KO – mKOk, LSS – LSSmOrange, S – mScarlet, PSS-Ponceau S staining.

For in-gel fluorescence analysis, 1 mL of each sample at mid-log phase was centrifuged (10 min at 2000 x *g*), the supernatant discarded, and the pellet was either snap-frozen in liquid nitrogen and stored at −80 °C for future analysis, or resuspended in ∼150–200 μL of sample buffer (0.2 M SDS, 0.3 M Tris, 30% glycerol, 0.02% bromophenol blue), vortexed and then incubated at 40 °C for 5 min on a heat block and quickly placed on ice until loaded on an SDS-PAGE gel. 15 μL of each sample was loaded along with 4 μL of PageRuler Plus Prestained Protein Ladder (Thermo Fisher Scientific, UK). After electrophoresis separation, gels were visualized in the ChemStudio PLUS using the filter combinations listed in Figure 3E. 504/10 nm excitation and no emission filter, was used to visualize the pre-stained protein ladder. Lastly, Coomassie Brilliant Blue (CBB) or Ponceau S staining of the protein gels was done to compare sample loading. All filter settings for the fluorescent screening of transformants on the Analytik Jena are provided in Supplemental File S5.

### 2.7. Flow cytometry

Flow cytometry of wild-type and fluorescent protein-expressing cells was performed using an Invitrogen Attune NxT flow cytometer (Thermo Fisher Scientific, UK) equipped with a 488 nm blue excitation laser for forward scatter (FSC), used to measure size, and with lasers for excitation at wavelengths 405 nm (Violet, VL), 488 nm (Blue, BL), 561 nm (Yellow, YL), and 638 nm (Red, RL), along with corresponding sets of emission filters supplied by the manufacturer. For dual population analysis, transformants and wild-type cells were grown in a 24-well plate as shown in Fig. 1b until they reached the mid-log phase. Each sample was diluted 1:100 with 0.9% (w/v) NaCl solution and 500 μL of each transformant and 500 μL of wild-type cells were combined in a 2 mL Eppendorf and measured through the sample injection port (100 μL sample/min). The first 15 μL of the combined sample was not recorded to ensure a stable cell flow rate during measurement. The Attune NxT Software v3.2.1 (Life Technologies, USA) was used to group cell populations and conduct further post-acquisition analyses.

### 2.8. Quantification of heterologous isoprene biosynthesis

To investigate the overexpression of the *I. batatas* isoprene synthase in *C. merolae* and the consequent production of isoprene, a transformant line was cultivated in 400 mL Algem photobioreactors (Algenuity, UK) along with the wild-type 10D and an mVenus-expressing strain as negative controls. Strains were cultivated under different light regimes at 42 °C and 120 rpm agitation. Preculturing before growth and isoprene analysis was performed by inoculation of cells to Algem flasks with MA2G medium for 96h, then, each strain was resuspended in the same medium to an OD_740nm_ ∼0.5 at the start of the experiment. Cell lines were grown in biological duplicates under constant illumination at 1200 μE or 750 μE, or under 12 h:12 h light: dark light cycling with the same light intensities.

Volatile isoprene was monitored during growth using a real-time headspace gas analysis system equipped with a triple filter mass spectrometer and multi-port inlet (Hiden Analytical HPR-20 R&D, UK). All cultures were continuously sparged with a 3% CO_2_ in air mix (25 mL/min), and the off-gas of each reactor was redirected through separate gas lines, first to a 250 mL bottle containing 80 g of CaCl_2_ used as a desiccant. Then, the dried gas was further directed to a 20-port inlet of the online headspace gas analysis system. The gas composition of each flask’s headspace was analyzed with a rotation to the next gas stream occurring every ∼3.5 mins. Isoprene was quantified by monitoring atomic mass units (amu) 67, 68, 53, 39, 40, 41, and 27; any overlapping amu of other gases present (O_2_, CO_2_, N_2_, Ar) were automatically deconvoluted by the Hiden Analytical QGA Software (version 2.0.8).

The amount of isoprene detected in the headspace of each culture was integrated over time, and the isoprene output (in ppm) reported by the instrument was corrected by comparison against a standard curve (Supplemental File S6) generated with flasks that were also kept in the Algems but contained different concentrations of loaded pure isoprene standard. In total, 10 control flasks were set up to cover five concentrations (10, 5, 2.5, 1.25, and 0.625 mg of isoprene) in duplicate. The temperature of the standard flasks was gradually increased from 20 °C to 38 °C over 1 h and then held for 10 h, during which time the isoprene standard in the headspace was quantified. The integrated isoprene amounts were plotted against the amounts added to each flask to generate the standard curve. The Hiden Analytical QGA Software was employed for real-time gas analysis until the end of algal cultivations (Figure 4E).

**Figure 4.**
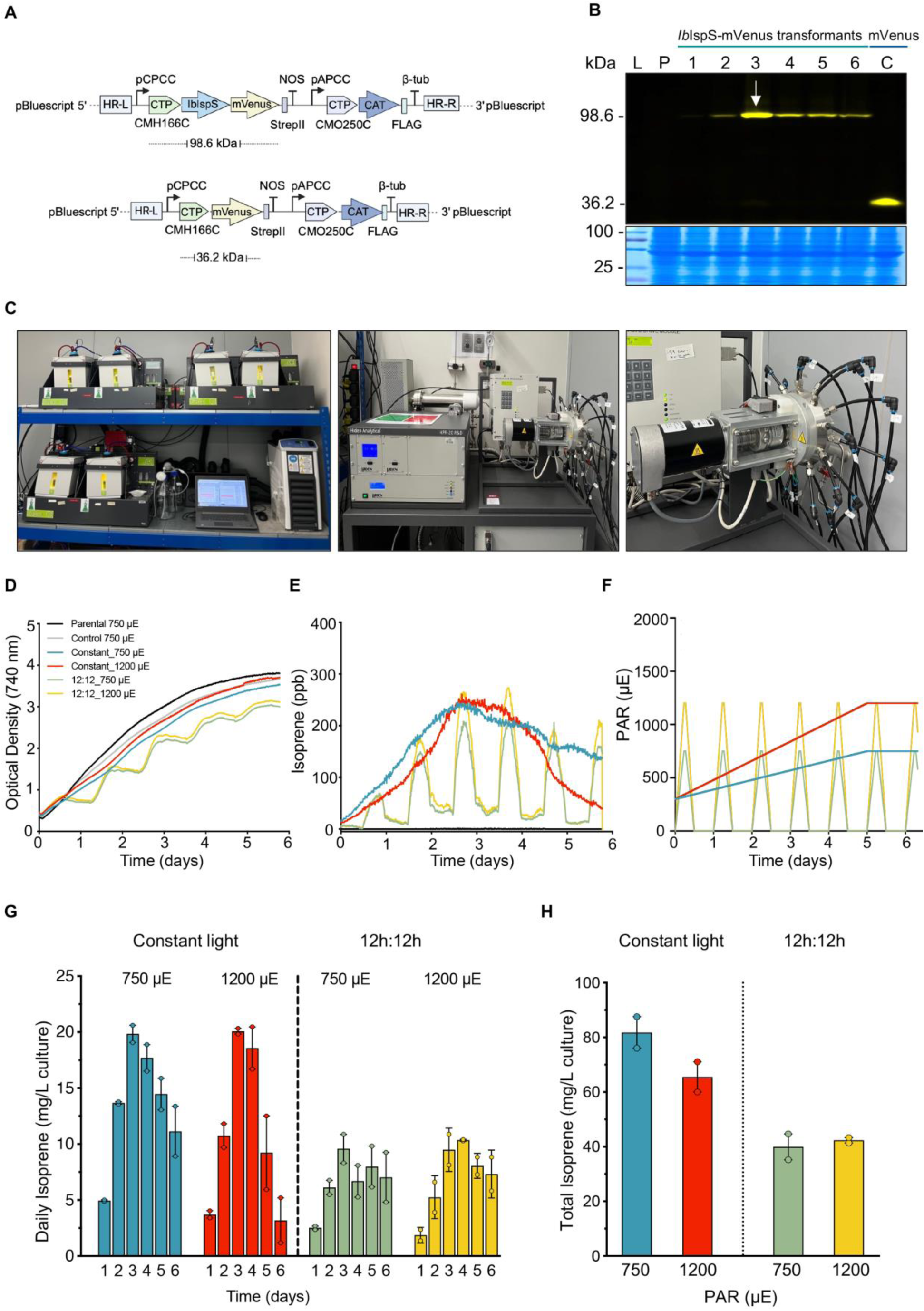
Characterization of isoprene production in an *Ib*IspS expressing *C. merolae* transformant. **A** – Isoprene synthase (IspS) expression plasmid (top) and negative control plasmid expressing mVenus (bottom). The *Ib*IspS gene was directly fused to mVenus and has a predicted molecular mass of 98.6 kDa in contrast to mVenus alone (36.2 kDa). **B** – Molecular mass of the direct fusion between *Ib*IspS-mVenus vs. mVenus alone confirmed by in-gel fluorescence analysis. Six *Ib*IspS-mVenus expressing transformants were compared and the transformant with the strongest expression (arrow) was selected for further experiments. **C** – from left to right: Algem photobioreactors (Algenuity, UK), Hiden Analytical HPR-20 R&D headspace gas analysis system (UK), and 20-port gas inlet. **D** – Optical densities at 740 nm (OD_740nm_) measured every 10 min during cultivation in Algem photobioreactors at different light intensities, and either constant light or under a 12 h light:12 h dark cycle. **E** – Volatile isoprene concentration in headspace under light regimes is shown in F. **F** – Light profiles used for each biological duplicate. For constant illumination, the light intensity increased linearly from 350 μE to the final experimental intensities (i.e., 720 and 1200 μE) after 120 h. **G** – Daily cumulative isoprene recorded in each condition. **H** – Cumulative isoprene produced during the entire cultivation period of 7 days.

### 2.9. Statistical analysis

Data analysis was conducted using both biological and technical replicates. Student *t*-test was carried out to determine if mean transformation efficiencies were significantly different (*p* < 0.05). Mean values and standard error (SEM) of replicates were calculated using the software JIMP Pro 16.2 (SAS Institute Inc., Cary, NC), and are illustrated in relevant graphs.

## 3. Results & Discussion

*C. merolae* 10D exhibits interesting properties like a small genome, ease of cell lysis, homologous recombination, and stable transgene expression that warrant its consideration as an emerging alga for biotechnology concepts. Here, we sought to develop a protocol that can more rapidly generate transgenic strains, characterize the behavior of integrated DNA, and present a useful molecular toolset for engineering its nuclear genome. We demonstrate the functionality of these tools in the proof-of-concept engineering of *C. merolae* 10D to produce volatile isoprene for the first time and characterize the dynamic nature of this production.

### 3.1. Development of a rapid chloramphenicol transformation protocol

A rapid transformation protocol for *C. merolae* 10D was developed, which generates single chloramphenicol-resistant colonies within 10 days (Figure 1). We employed a two-expression-cassette system which we designed *de novo* for this study (Figure 1A). The *APCC* and *CPCC* promoters were used to drive transgene expression as they produce high transcriptional activity under illuminated conditions (Fujiwara et al., 2019). *CPCC* was chosen for the gene of interest cassette as this promoter exhibited higher recombinant protein accumulation than the *APCC* in a previous study (Fujiwara et al., 2019). The *APCC* promoter was used to express the selectable marker CAT. Homologous recombination for transgene integration was directed by 550 bp sequences (Figure 2A,B) targeting the CMD184C–185C locus found on *C. merolae* 10D chromosome 4 (Fujiwara et al., 2017).

The gene expression cassettes were amplified by PCR and used in the PEG-mediated transformation of *C. merolae* 10D (Figure 1A-B). The alga can take up foreign DNA without a physical agitation agent like the glass beads used for cell wall deficient *Chlamydomonas reinhardtii* (Kindle, 1990). Salmon sperm DNA was used in the transformation mix along with 1 pmol of linearized transgene cassette DNA. Salmon sperm DNA is denatured before use and is typically employed as a DNA carrier to improve transformation efficiency in yeast (Longmuir et al., 2019) and marine microalgae (Zhang & Hu, 2014). The recovery of transformants as single colonies requires selection on 20% cornstarch slurry spots on freshly prepared chloramphenicol-containing MA2-G plates.

Compared to the top-starch method (Takemura, Kobayashi, et al., 2019), this is a more labor-intensive protocol, as the plates must be prepared fresh; cornstarch slurry spotted, and cells serially diluted before inoculation. However, the use of a 3X serial dilution of cells before plating allows for balancing the culture density to enable single colonies to appear without overcrowding. Once colonies are visible, they can be directly picked into liquid MA2G medium to later inoculate a maintenance starch spot for long-term storage, and the liquid culture can be used directly for further scale-up and screening analysis (Figure 1B). Following this new experimental pipeline, the time required to generate transformants is reasonable for conducting iterative engineering cycles required for most biotechnology concepts.

Both linear and circular plasmid constructs containing mVenus were transformed into *C. merolae* 10D to determine differences in transformation efficiencies and potential transgene expression differences. It has been recently shown in the red alga *Porphyridium purpureum* that the bacterial origin of replication could confer autonomous replication of circular, episomal, plasmids within cells and consequent higher recombinant protein titers (Hammel et al., 2024; Li & Bock, 2018). The efficiency of transforming PCR linearized constructs in *C. merolae* 10D was 18-fold higher than in the episomal transformation (*p* < 0.0001; Figure 1E). Episomal transformation was carried out with the same workflow as linear, but the plasmid was not linearized before the transformation. The number of colony-forming units in the linear and circular transformation plates was counted 10 days after plating. Only a few colonies were recovered when the cells were transformed with a circular plasmid.

Efficient gene targeting and stable transgene expression represent two advantages of *C. merolae* 10D as a host for synthetic biology and metabolic engineering. The intergenic region within the CMD184C-CMD185C loci was selected as the target site, as it has been previously used successfully (Fujiwara et al., 2013, 2017; Fujiwara & Ohnuma, 2017). The occurrence of HR at this locus was examined by PCR. Integration of the target binary cassette carrying mVenus and CAT was observed in 22 out of 24 transformants (92%; Figure 2C). Previously, targeted integration of the CAT selection marker was observed in 24 out of 33 transformants (73%), when the transgene was flanked with 500-bp homology arms and the cells were selected in a liquid medium with 200 μg/mL chloramphenicol for 10 days and then plated for another 2 weeks (Fujiwara et al., 2017). In our efforts to obtain transformants more rapidly, direct plating on 250 μg/mL chloramphenicol after 48 h recovery was employed. This modification was able to speed the CAT transformation up by two weeks. Two transformants (7 and 8) presented the same amplicon size as the wild-type (∼ 1 kb) indicating integration outside the intended region. Although not directly integrated at the target locus, these transformants exhibited mVenus fluorescence (Figure 2C). Both at the agar plate and in-gel fluorescence levels, the mVenus reporter was detected in all 24 transformants. Across the clones, a consistent mVenus signal was observed, except for those with non-targeted integration (7 and 8) and clones 20 and 21 (Figure 2C). These transformants, as well as properly-integrated and moderate mVenus-expressing colonies 4, 5, 23, and 24, were selected for assessment of gene copy number by qPCR. All clones with higher fluorescence than the average exhibited 2 or more copies of the transgene construct. Colonies 8 and 20, with the highest mVenus fluorescence, appear to have 30 and 20 copies of the transgene in their genomes respectively. This is similar to a previous report of multicopy insertion wherein tandemly repeated insertion of the introduced fragments was observed (Fujiwara et al., 2013). A semi-quantitative assessment of mVenus accumulation was also carried out in these eight transformants. The TSP fraction of the transformants was quantified by the BCA test and compared against a dilution series of purified mVenus using an SDS-PAGE gel and fluorescence imaging. Recombinant protein levels accumulated to between 0.3-0.6% of total soluble protein (Figure 2E). The copy number of gene insertion likely influences the levels of recombinant protein within the cell but does not match the levels reported for *P. purpureum* episomal transgene expression (i.e. up to 5% of the total soluble protein (Li & Bock, 2018)).

### 3.2 A suite of new fluorescent reporters for *C. merolae*

New fluorescent reporters applied in *C. merolae* can enable combinations for protein interaction studies, or as fusion partners to target recombinant proteins. We designed codon-optimized transgenes for the expression of fluorescent reporters mTagBFP2, mTFP1, Clover, mVenus, mKOk, LSSmOrange, and mScarlet (Figure 3A-B). mVenus and mScarlet have been previously used in this host (Fujiwara et al., 2021; Seger et al., 2023), however, the addition of broader spectral tools can expand applications in this host. Here, stable expression of these fluorescent reporters was demonstrated using the same transformation and recovery protocols as outlined above (Figure 3C-E). Cell populations for each reporter could be distinguished from that of the wild-type cells in flow cytometry analyses and represent a sensitive analytic technique where fluorophores could be multiplexed (Figure 3C). Additionally, visualization of transformants and their expression at the agar plate level and in protein gels can aid gene expression assessment and transformant status. We compared *C. merolae* with our previous efforts in practical fluorescence imaging developed for the green alga *C. reinhardtii* (Figure 3D-E, as reported in (Gutiérrez et al., 2022)). We applied different bandpass filters in a modified gel doc system to visualize pre-selected transformants either in agar plates or protein gels and show that their signal can be visualized separately from chlorophyll *a* and phycocyanin autofluorescence (Gutiérrez et al., 2022). The emissions of mTagBFP2, mVenus, mKOk, and LSSmOrange were visible without spectral overlap, and to a lesser extent mTFP1, mVenus, and mScarlet could be combined in this analysis (Figure 3D-E and Supplemental File S5). The ability to conduct targeted HR into the *C. merolae* genome means fewer transformants must be screened compared to efforts in green algae (Abdallah et al., 2022; Gutiérrez et al., 2024) and the ability to rapidly detect reporter expression is a practical asset for analyzing trangene expression.

### 3.3 Overexpression of *Ib*IspS results in volatile isoprene production

We wanted to test the heterologous expression of isoprene synthase in *C. merolae* using the transformation and some of the screening tools described before. Terpenoids, also known as isoprenoids, constitute a diverse class of natural compounds many of which are essential metabolites found in all organisms where they play roles in regulating electron transfer, participate in photosynthesis, and a host of other cellular functions (Masyita et al., 2022). Isoprenoids are generated from either the 2-C-methyl-D-erythritol 4-phosphate (MEP) or mevalonate (MVA) pathways to produce the 5-carbon prenylated precursors of terpenoids, isopentyl- or its isomer dimethylallyl-diphosphate (IPP and DMAPP, respectively) (Lichtenthaler, 1999; Perez-Gil et al., 2024; Pu et al., 2021). DMAPP is used by many plants to reduce reactive oxygen species stress through the cleavage of the diphosphate group and release of the 5-carbon volatile molecule isoprene through the action of isoprene synthases. Isoprene production has been engineered into several microbes, both as a means by which to determine how much carbon flux can be diverted to this product and because it represents a bulk platform chemical for use in the production of rubber or jet fuels (Aldridge et al., 2021; Chaves & Melis, 2018; Diner et al., 2018; Gomaa et al., 2017; Rana et al., 2022; Yahya et al., 2023). Based on in *silico* studies, only the MEP pathway is found in the chloroplast of *C. merolae,* where it generates all required precursors for cellular terpenoids (Grauvogel & Petersen, 2007; Lohr et al., 2012). *C. merolae* does not naturally produce volatile isoprene and we found no evidence of isoprene synthase in its genome. We sought to determine whether we could engineer the production of isoprene from *C. merolae* using our modular plasmid toolkit. We chose this volatile product as the alga can grow to high densities on CO_2_ and its cultivation temperature (42 °C) is higher than the boiling point of isoprene (34 °C) which should facilitate its production.

Many isoprene synthases (IspSs) have been heterologously expressed in microbial hosts, including IspS from the poplar tree (*Populus alba*; (Gao et al., 2016)), the Kudzu vine (*Pueraria montana; (Gomaa et al., 2017)*), and eucalyptus (*Eucalyptus globulus*; (Englund et al., 2018; Gao et al., 2016)). The IspS gene from *I. batatas* (sweet potato; *Ib*IspS) was chosen for heterologous expression in *C. merolae* as it exhibited robust isoprene production in our previous tests in the green alga *C. reinhardtii* (Yahya et al., 2023). The amino acid sequence for *Ib*IspS was synthetically optimized for expression from the *C. merolae* nuclear genome after removal of its native plastid targeting peptide and expressed as an N-terminal fusion with mVenus (Figure 4A). This was done both to facilitate the identification of robustly expressing clones and to aid protein stability (Grauvogel & Petersen, 2007; Lohr et al., 2012; Yahya et al., 2023). The native plastid transit peptide from DNA Gyrase B was used to target the heterologous enzyme to the algal plastid. We also chose this enzyme as an *in vitro* study had shown the optimum catalytic activity for this enzyme was 42 °C, the same as the alga’s optimum cultivation temperature (Li et al., 2019), adding to our interest in its use in *C. merolae*.

After transformation of the linearized plasmid, in-gel fluorescence screening was used to identify the fusion protein expression in transformants selected for mVenus signal. The correct molecular mass of the fusion protein was observed (Figure 4B). The transformant showing the highest expression was then chosen and grown in 400 mL Algem photobioreactors (Algenuity, UK) to test growth and isoprene accumulation over 6 days using different light regimes in parallel photobioreactors using a multi-port online headspace analyzer (Figure 4C–H). The parental *C. merolae* 10D, as well as an mVenus expressing control, were also grown in parallel and their headspaces were compared to the transformant. The transformant was subjected to either day-night cycling or 24-hour continuous illumination using two different settings.

All strains grew in their respective conditions as observed by live monitoring of optical density throughout the cultivation (Figure 4D). No isoprene was detected in either parental or mVenus alone controls, however, isoprene was continuously detected from the transformant expressing the *Ib*IspS (Figure 4E). The isoprene detected was dependent on illumination conditions, with both continuous illumination cultures reaching the same maximum output by day 3 and exhibiting a reduction in the later stages of cultivation (Figure 4E). This did not coincide with the optical density measurements, which continued to increase until day 5 (Figure 4D). The cultures with 12:12 illumination and dark cycles exhibited daily isoprene production which followed the increasing and lowering of the light programs (Figure 4F). This may indicate that the *CPCC* promoter is dynamically controlled by light as very low levels of isoprene were observed in non-illuminated conditions daily. Isoprene production was lower in constant 1200 µE illumination than 750 µE, which may indicate other levels of pathway flux regulation if the promoter is light-controlled. The daily accumulated isoprene production was measured (Figure 4G) and totaled (Figure 4H), with the overall highest amount in the continuous illumination at 750 µE light yielding 80 mg isoprene/L culture in 6 days. This is already higher than ∼55 mg/L achieved by expression of the same gene alone in the green alga *C. reinhardtii* (Yahya et al., 2023). The productivity was highest for both continuous cultivations on days 3-4 (Figure 4G), which may also mean growth stage and culture densities play a role in transgene expression. Further systematic investigations of promoter expression rates during cultivation and influences of cultivation conditions will be the subject of follow-up studies.

## Conclusions

In this work, we optimized the CAT transformation and selection protocol. This reduces the time from transformation to colony selection from about four weeks to 10 days. This shortened time frame will be beneficial for the often necessary iterative transformations involved in engineering strains. *C. merolae* 10D is a promising alternative photosynthetic cell chassis to green algae because of its unique cultivation conditions, possibilities of transgene HR integration, and stable transgene expression. In this work, we used *in-silico* design and *de novo* construction of transgene expression constructs to demonstrate the speed with which engineering concepts can be implemented in this host. Through this optimized transformation and selection protocol, colonies can be recovered within a reasonable time frame to enable iterative engineering activities. We demonstrated the production of engineered volatile isoprene as a proof of concept, but many other targets that require the introduction of multiple gene pathways are now more feasible to investigate. Given its stable cultivation and requirement for photoautotrophic growth, *C. merolae* is an interesting algal host for scaled cultivation and bioproduction concepts. Further understanding of the behavior of promoters and terminators under different conditions will be required to determine how expressed genes will behave in different cultivation conditions. *C. merolae* represents an interesting emerging model alga that could be used for many biotechnological investigations. Future work should broaden its molecular toolkit with antibiotic-based, counter-selectable markers and further examples of metabolic engineering for which the work presented here will be a useful foundation.

## Supporting information

Supp File S1 - Data Table Sequence information

Suppl File S2 - Full annotated plasmids

Suppl File S3 - Transformation efficiencies

Suppl File S4 - PCR primers and settings

Suppl File S5 - Filter settings Analityk Jena

Suppl File S6 - Isoprene culture

Suppl Table 1 - Plasmid numbers

## Acknowledgments

The authors acknowledge the labs of Shin-Ya Miyagashima and Kan Tanaka for foundational studies of *C. merolae*. KJL acknowledges baseline research funding from KAUST. MVV and MLPM are funded by the KAUST opportunity fund grant #5576 awarded to KJL. SO is funded by a corporate research project with the Saudi International Petrochemical Company (Sipchem) awarded to KJL. PJL acknowledges funding from Xylem, Inc. and ASU Lightworks. SDR and MRS acknowledge funding from the NSERC Discovery Grant program and UNBC’s Office of Research and Innovation.

Conflict of interest

The authors declare that they have no conflict of interest.

## Supplementary figures legends

**Suppl. Figure 1.** Full-length images of the molecular screening of mVenus-expressing transformants. **A** – Polymerase chain reaction (PCR) to confirm integration of the linearized plasmid at the D184-185 neutral locus (insert) and presence of the selectable marker (CAT) in 24 transformants. **B** – Plate-level fluorescence analysis of the same transformants on Gellan gum plates stained with amido black. 10 μL of liquid culture were spotted per transformant along with the parental strain (top) to compare for Chlorophyll *a* vs mVenus fluorescent emission, measured at the indicated wavelengths using ChemStudio Plus. **C** – In-gel fluorescence also used to assess the expression level of mVenus in the 24 transformants. The expected molecular weight of mVenus is ∼26 kDa. CBB: Coomassie brilliant blue stain included as a loading control.

**Suppl. Figure 2. A-F** – Raw in-gel fluorescence images captured using different filter sets. Filters and exposure times for different fluorophores are indicated. Images were captured in ChemStudio plus. **G** – Ponceau S staining is also included as a loading control.

**Supp. Figure 3.** Quantitative PCR (qPCR) optimization **A** – Electrophoresis gel showing the results from a thermal gradient PCR to determine the optimal annealing temperature of the primers. A no-template control (NTC; DNA replaced with water) was included. Arrows indicate the best-predicted temperature (∼60 °C) for both primer sets targeting regions within mVenus and 60S rDNA (reference gene). **B** – Standard curve to measure the efficiency of the amplicon doubling. 91.0% and 109.6% were obtained for 60S rDNA and mVenus, respectively. **C-D** – Template amplification for both target genes when 5 concentrations of genomic DNA were tested (1/5, 1/25, 1/125, 1/625, and 1/3125). **E-F** – Meltcurve generated by the CFX Maestro software. One peak confirms one product is being generated without the presence of non-specific binding. **G** – Cq values of the amplification of 60S rDNA and mVenus for 8 different transformants. Cq values were measured in technical triplicates for each primer set and the standard error is expressed with the error bars.

## Abbreviations

BCA: Bicinchoninic Acid

CAT: Chloramphenicol transferase

CBB: Coomassie Brilliant Blue

cfu: Colony-forming units

CTP: Chloroplast targeting peptide

HR: Homologous recombination

IspS: Isoprene synthase

NTC: No template controls

PEG: Polyethylene glycol

qPCR: Quantitative PCR

SDS-PAGE: Sodium dodecyl sulfate-polyacrylamide gel electrophoresis

TSP: Total soluble protein

