## Supplemental Figures 1-3 for "A rapid CAT transformation protocol and nuclear transgene expression tools for metabolic engineering in *Cyanidioschyzon merolae* 10D"

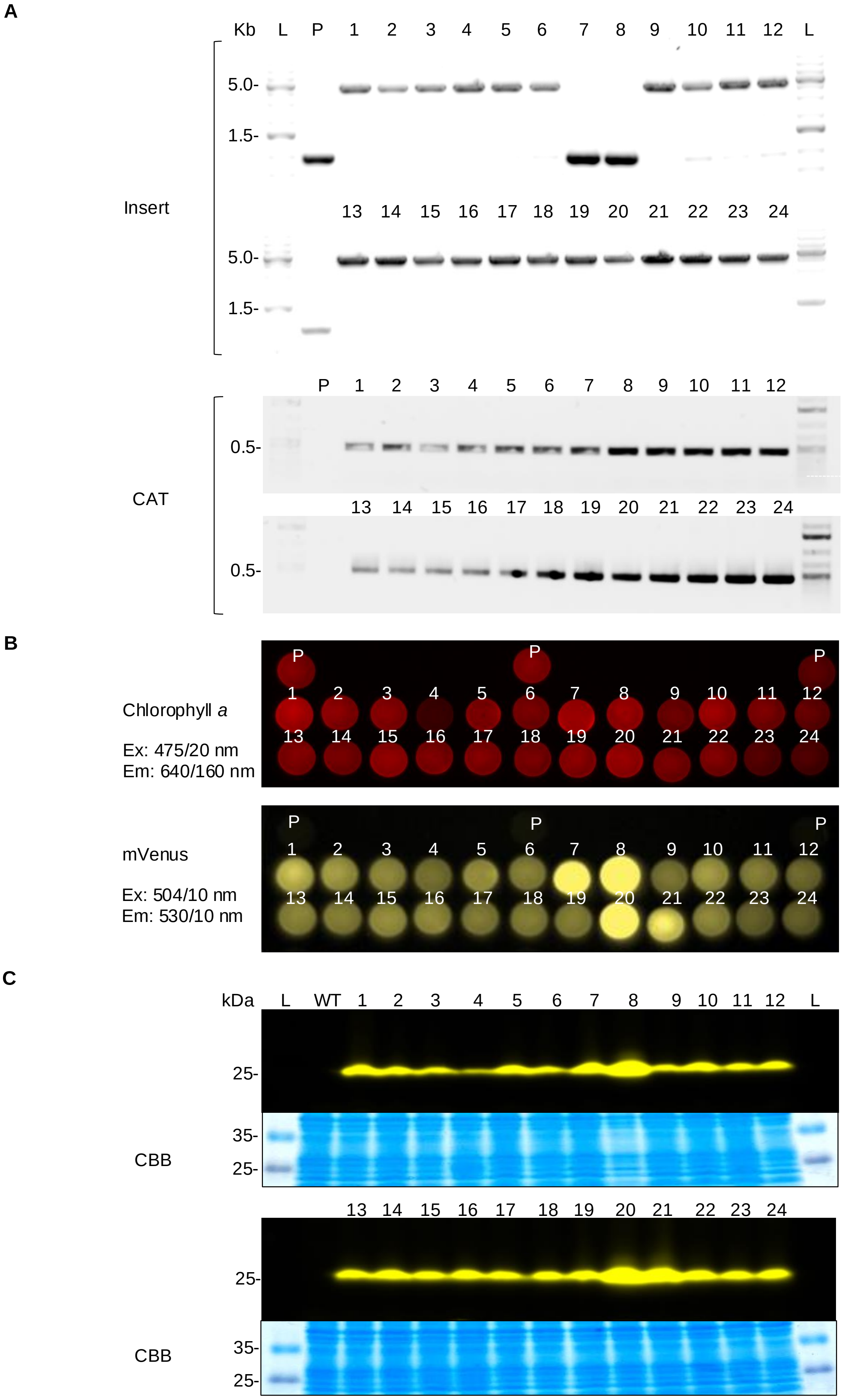


**Suppl. Figure 1.** Full-length images of the molecular screening of mVenus-expressing transformants. **A** – Polymerase chain reaction (PCR) to confirm integration of the linearized plasmid at the D184-185 neutral locus (insert) and presence of the selectable marker (CAT) in 24 transformants. **B** – Plate-level fluorescence analysis of the same transformants on Gellan gum plates stained with amido black. 10 µL of liquid culture were spotted per transformant along with the parental strain (top) to compare for Chlorophyll *a* vs mVenus fluorescent emission, measured at the indicated wavelengths using ChemStudio Plus. **C** – In-gel fluorescence also used to assess the expression level of mVenus in the 24 transformants. The expected molecular weight of mVenus is ~26 kDa. CBB: Coomassie brilliant blue stain included as a loading control.

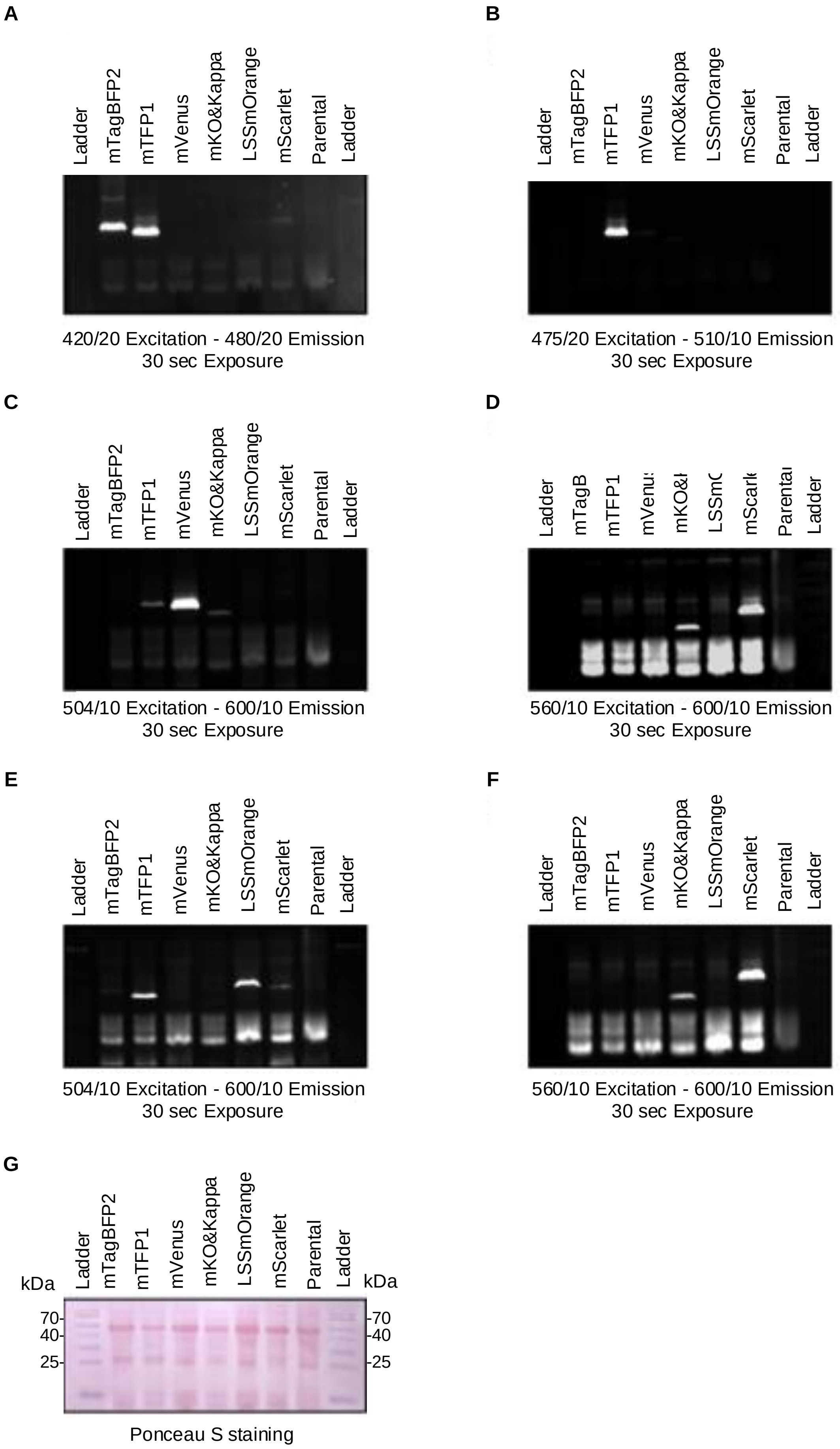


**Suppl. Figure 2. A-F** – Raw in-gel fluorescence images captured using different filter sets. Filters and exposure times for different fluorophores are indicated. Images were captured in ChemStudio plus. **G** – Ponceau S staining is also included as a loading control.

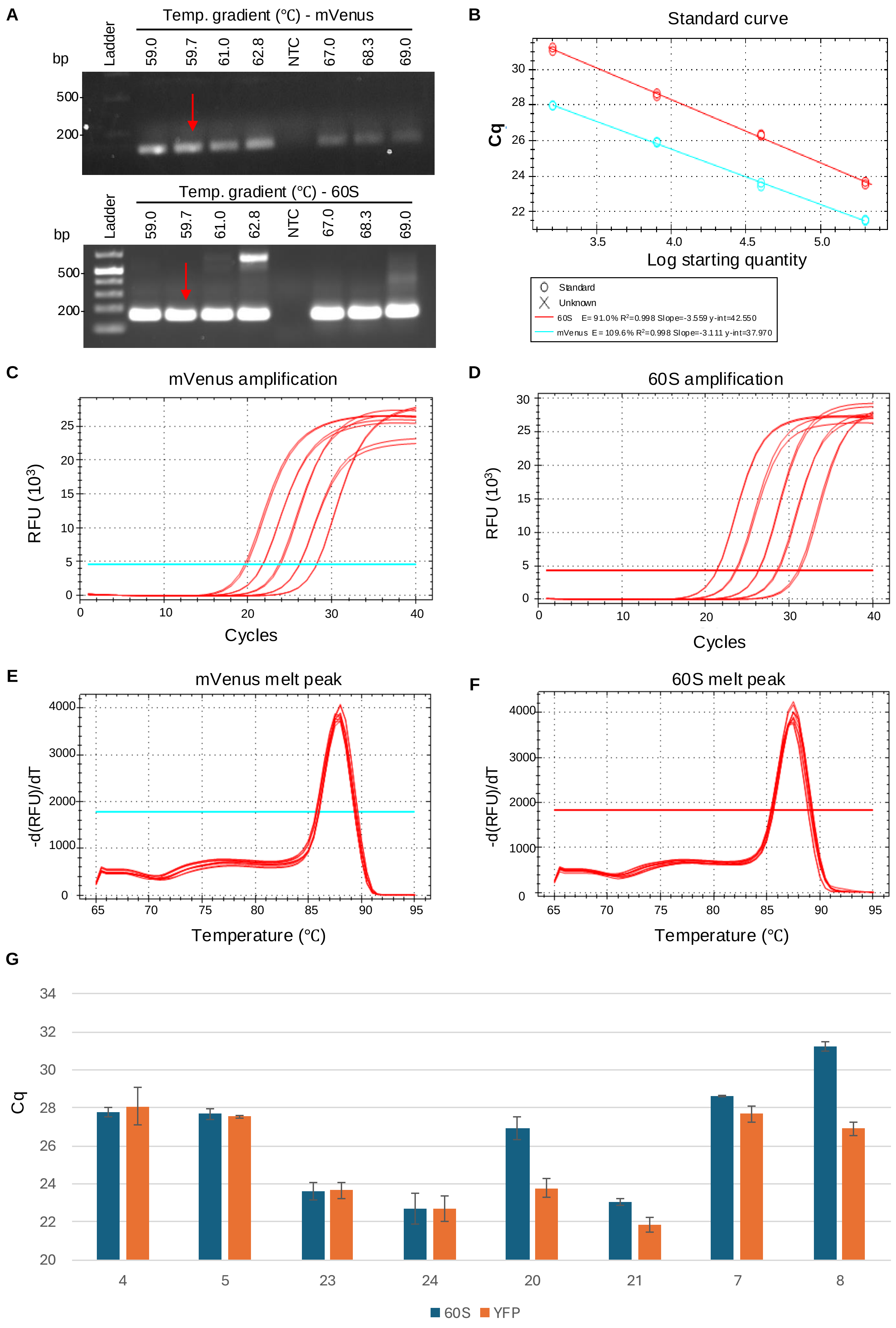


**Supp. Figure 3.** Quantitative PCR (qPCR) optimization **A** – Electrophoresis gel showing the results from a thermal gradient PCR to determine the optimal annealing temperature of the primers. A no-template control (NTC; DNA replaced with water) was included. Arrows indicate the best-predicted temperature (~60 ºC ) for both primer sets targeting regions within mVenus and 60S rDNA (reference gene). **B** – Standard curve to measure the efficiency of the amplicon doubling. 91.0% and 109.6% were obtained for 60S rDNA and mVenus, respectively. **C-D** – Template amplification for both target genes when 5 concentrations of genomic DNA were tested (1/5, 1/25, 1/125, 1/625, and 1/3125). **E-F** – Meltcurve generated by the CFX Maestro software. One peak confirms one product is being generated without the presence of non-specific binding. **G** – Cq values of the amplification of 60S rDNA and mVenus for 8 different transformants. Cq values were measured in technical triplicates for each primer set and the standard error is expressed with the error bars.
